# The Hidden Cost of Blue Light: Multiorgan Pathology Induced by LED Exposure in Mice

**DOI:** 10.64898/2026.09.11.750850

**Authors:** Heba Al-Hussaini, Leora D’Souza, Sonia M. Hasan, Sampath Madhyastha, Mohammed Al-Onaizi, Sakkattu Rao, Rawan Kittaneh, Batool Samaro, Bara S. Abed, Glen Jeffery

## Abstract

Life evolved under broad-spectrum sunlight (300–3500 nm), but most modern light-emitting diodes (LEDs) emit within a restricted visible range (350-650 nm) leaving lighting environments that are short-wavelength dominant. We ask if chronic 450nm exposure dominant in LEDs alters metabolism and organ integrity in ageing mice environmentally exposed at 13 mW/cm² for 5 h/day over 40 weeks, at an intensity within the range of human environmental exposures. Mice exhibited progressive weight gain despite reduced food intake, with altered glucose tolerance and insulin sensitivity over time. White adipose tissue mass increased, while brown adipose tissue weight and UCP1 protein levels remained unchanged. Relative organ weights (heart, liver, kidney, testis) were each reduced by ∼10%, with histopathology revealing fat deposition, hepatocellular degeneration, renal tubular damage, steatosis, and impaired testicular architecture. These findings demonstrate that chronic 450 nm light exposure is associated with metabolic alterations and multiorgan pathology in mice.

## Introduction

Solar light has a wide spectral range from the ultraviolet to the infrared (300-3500 nm) that has been consistent for billions of years. Short wavelength elements in this light impact metabolism by modulating mitochondrial function, an effect that is conserved across species^1–4^. Longer wavelengths (∼650 nm and above) that penetrate tissue deeply^4^ enhance mitochondrial performance, improving cellular respiration, particularly when it has declined due to age and disease. These wavelengths increase mitochondrial membrane potential and production of adenosine triphosphate (ATP)^5^. They also positively influence health span in shorter-lived animals^5^. In humans and other animals, they improve age-related visual function^3,4,6^. Longer wavelengths also have the ability to regulate blood sugars by increasing mitochondrial demand for carbohydrates^1, 2^. Conversely, some shorter wavelengths (400-450 nm) reduce the same metrics and are associated with shorter health spans^7, 8^. They also have the ability to rapidly shift heart rate and blood pressure^9^.

In sunlight, these different parts of the spectrum are in balance. However, lighting in the built environment has recently shifted from a wide sunlight spectrum, which is also found in incandescent luminaries, to the restrictive, blue-dominant spectrum of LEDs (400-650 nm), which lacks longer wavelengths and often have a peak in the 420-450 nm range. The significance of the negative impact of LED lighting is revealed in humans when short-term incandescent (400-1500 nm) supplementation is given in LED illuminated environments. This significantly improved colour contrast sensitivity, with effects persisting for up to two months ^10^.

We have previously shown that mice exposed to LED lighting in the 420-450 nm range that reduces mitochondrial performance, suffer from increased weight gain, disrupted serum cytokine patterns, and abnormal explorative behaviour^11^. These changes appear systemic and consequently may impact more widely. Prolonged short-wavelength exposure may influence physiological systems beyond the retina and central nervous system. Although previous studies have primarily focused on retinal, circadian, behavioural, or metabolic responses to short wavelength light, the long-term effects of chronic 450-nm exposure on multiple peripheral organs remain poorly characterized. Here we explore this suggestion, examining an array of organs in mice exposed to 450 nm light at irradiance levels within the range of human exposure in the built environment. The results reveal widespread pathological features in each of the organs examined and raise concerns regarding public health in the built environment where people spend most of their time.

## Results

### Light penetration

No detectable transmission of 450 nm light was observed through excised mouse skin when measured with a radiometer and spectrometer, indicating that this wavelength does not penetrate beyond the superficial tissue layers. Fur may have been an additional barrier. This confirms that internal organs were not directly exposed to 450 nm light during the experiment. Therefore, the systemic metabolic and multiorgan changes observed in experimental mice are likely mediated through indirect physiological mechanisms.

### Body weight and food intake

Mice maintained under 450 nm LED lighting exhibited a progressive increase in body weight compared to controls (Fig 1), consistent with our previous findings^11^. Differences became statistically significant (p<0.001) by week 10 and remained significant for the duration of the study. Notably, this weight gain occurred despite the experimental group consuming approximately 10–15% less food than control animals (p<0.05).

**Figure 1.**
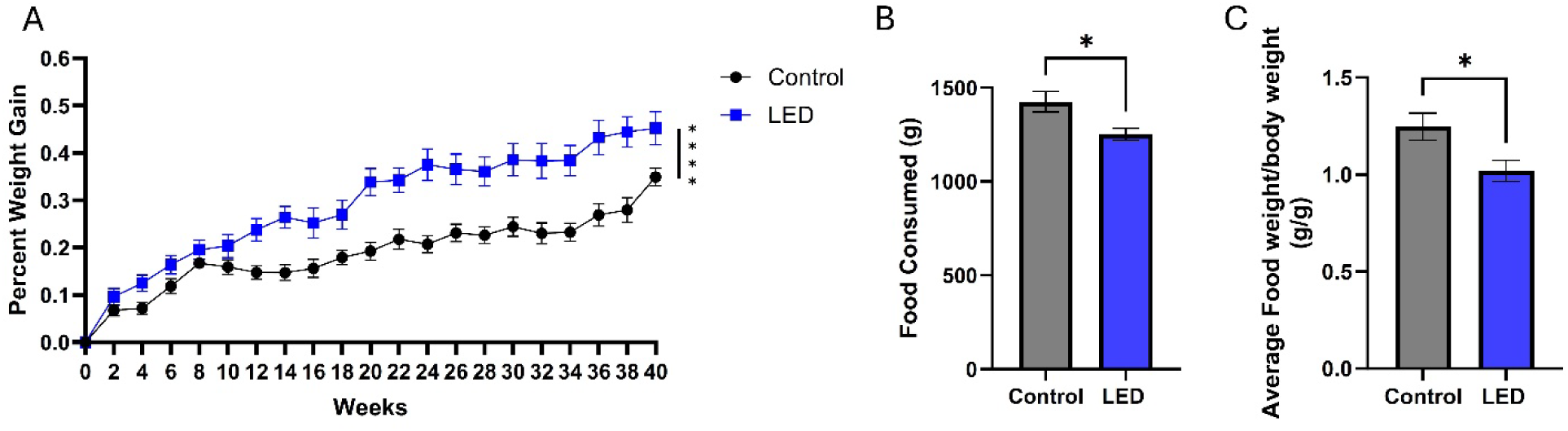
Body weight. Blue light of wavelength 450 nm causes weight gain and reduction in food intake (A) Percentage of weight gain in mice exposed to normal light or LED 450 nm over 40 weeks. Mice exposed to 450 nm light have significant weight gain from around 8-10 weeks of exposure. (B) Total food consumption (g) and (C) average weekly food consumption normalised to body weight (g/g) in control and LED exposed mice. Mice exposed to 420nm have a significant reduction in food intake in spite of increased weight gain. This is consistent with a metabolic shift. Error bars indicate SEM. *p<0.05, ****p<0.0001, n=10 per group. Data was analysed using two-way ANOVA for (A) and unpaired t-test for (B) and (C).

### Fat deposition

Chronic exposure to 450 nm LED light was associated with increased adiposity. Relative to body weight, both subcutaneous white adipose tissue (sWAT) and epididymal white adipose tissue (eWAT) weights were significantly higher in LED exposed mice compared with controls (p < 0.05 and p < 0.005, respectively, Fig 2A&B). In contrast, brown adipose tissue (BAT) weight did not differ significantly between the groups (Fig 2.C). Gross examination at necropsy confirmed greater accumulation of white adipose tissue in LED-exposed animals Fig 2.D&E). Assessment of BAT thermogenic markers showed no significant difference in UCP1 protein expression between control and LED-exposed mice (Fig 2. F). Together, these findings indicate increased white adipose tissue accumulation following chronic 450 nm LED exposure, without detectable changes in BAT mass or UCP1 expression (Fig 2. F-G).

**Figure 2.**
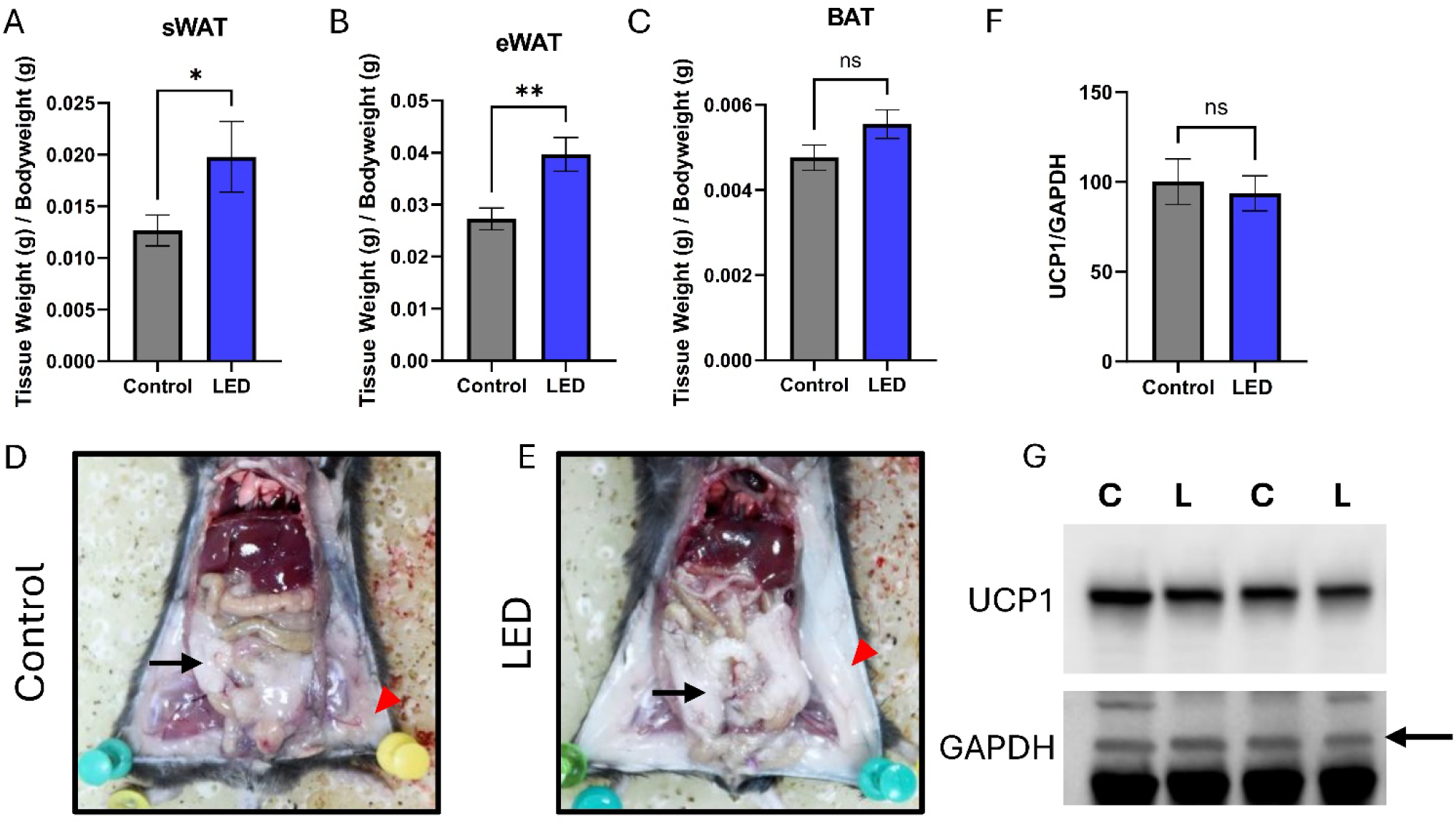
Fat deposition in mice exposed to 450 nm LED light. Tissue weight were normalized to body weight (g/g) for (A) subcutaneous white adipose tissue (sWAT), (B) epididymal white adipose tissue (eWAT), and (C) brown adipose tissue (BAT). Representative images showing sWAT (red arrowheads) and eWAT (black arrows) in mice exposed to (D) normal light and (E) 450 nm blue LED light. (F and G) Western blot analysis for UCP1 in brown adipose tissue showed no significant changes. Arrow indicates band for GAPDH. Error bars indicate SEM. *p<0.05, **p<0.01, n=10 per group for (A-C) and n=6 per group for (F-G). Data was analysed using unpaired t-test. Only representative bands are shown in panel (G). Original full length blots for (G) are presented in Supplementary Figure 1

### Metabolic results

At the start of the experiments and at four weeks of light exposure, no significant differences were observed between the control and LED groups in oral glucose tolerance, insulin tolerance, or fasting blood glucose levels (Fig 3. A-C). After 8 weeks of exposure, LED exposed mice exhibited significantly impaired oral glucose tolerance compared with control mice (Fig 3. A). This difference was most pronounced at Week 20 (p < 0.0001) and remained significant at Weeks 32 and 40. A similar pattern was observed during insulin tolerance testing. No differences were detected at baseline or at 4 weeks of exposure (Fig 3. B). However, while LED exposed mice showed a significantly altered response to insulin administration at Weeks 8, 20, and 32 compared with controls (Fig 3. B) at Week 40 there was no statistical difference between the two groups (Fig 3. B). A transient increase in fasting blood glucose was observed in the LED exposed group at Week 8 (p<0.01, Fig 3. C). However, there were no significant differences observed between groups at other recorded time points (Fig 3. C). Together, these findings demonstrate that chronic 450 nm LED exposure progressively impaired glucose and insulin tolerance while having a clear effect on fasting blood glucose levels only at 8 weeks.

**Figure 3.**
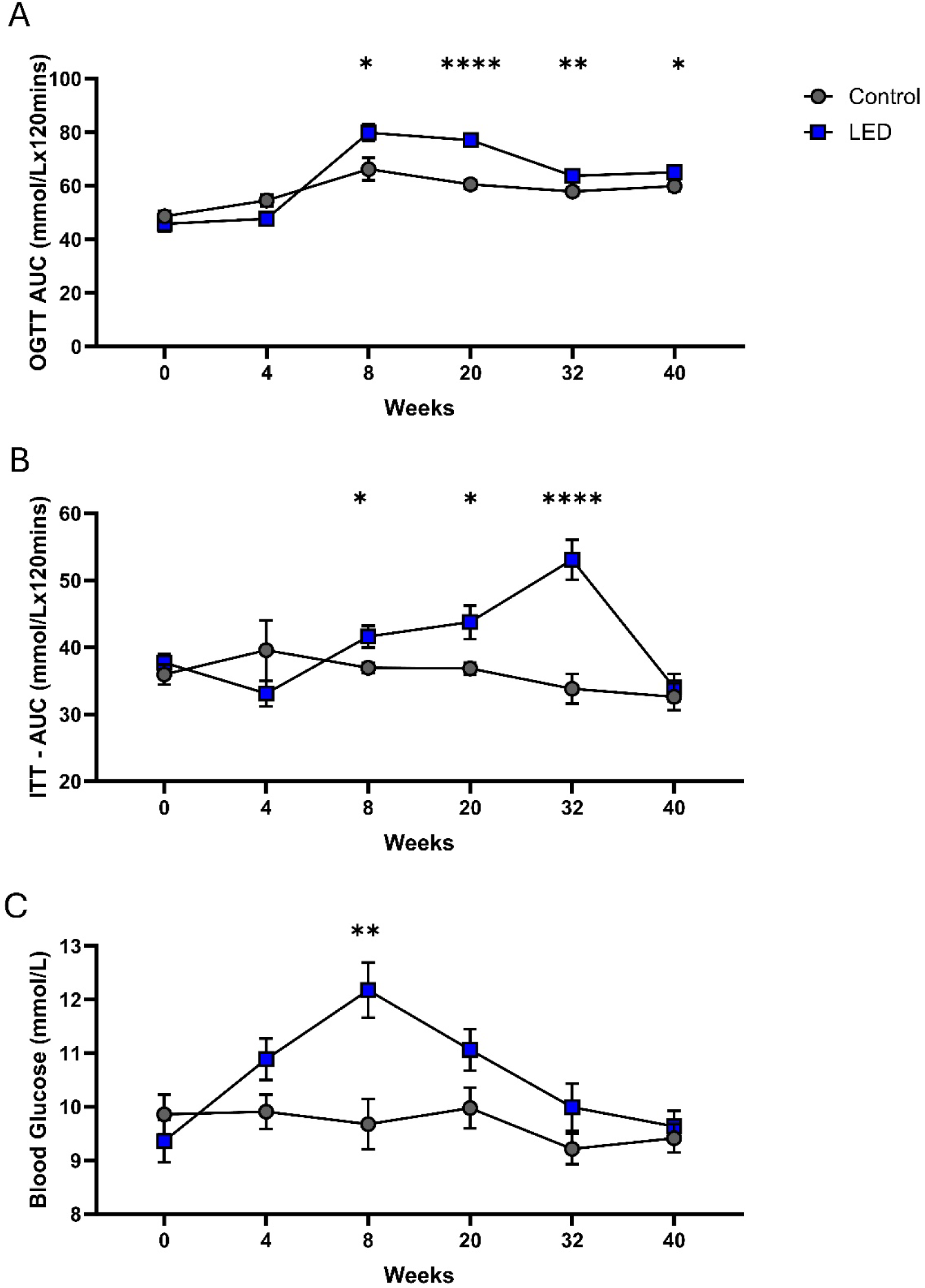
Disruption of Glucose Homeostasis. Mice exposed to 450 nm light develop glucose intolerance and insulin resistance. Data are shown for baseline (week 0), and at 4, 8, 20,32 and 40 weeks after treatment. Area Under the Curve (AUC) analysis for (A) oral glucose tolerance test and (B) insulin tolerance test. (C) Fasting glucose levels (mmol/L) for the same time points. Changes over the 40-week period are varied but indicate complex shifts in metabolism under 450 nm. Error bars indicate SEM. ns=not significant *p<0.05, **p<0.01, ****p<0.0001 n=10 per group. Data was analysed by unpaired t-test.

### Effect on multiple organs

The weights of key organs were reduced by about 10% in the mice exposed to 450 nm light compared to controls. In each case these differences were statistically significant: heart (p<0.01), liver (p<0.05), kidneys (p<0.01), and testes (p<0.05) (Figs 4. A-B, 5. A, 7 and 9. A).

**Figure 4.**
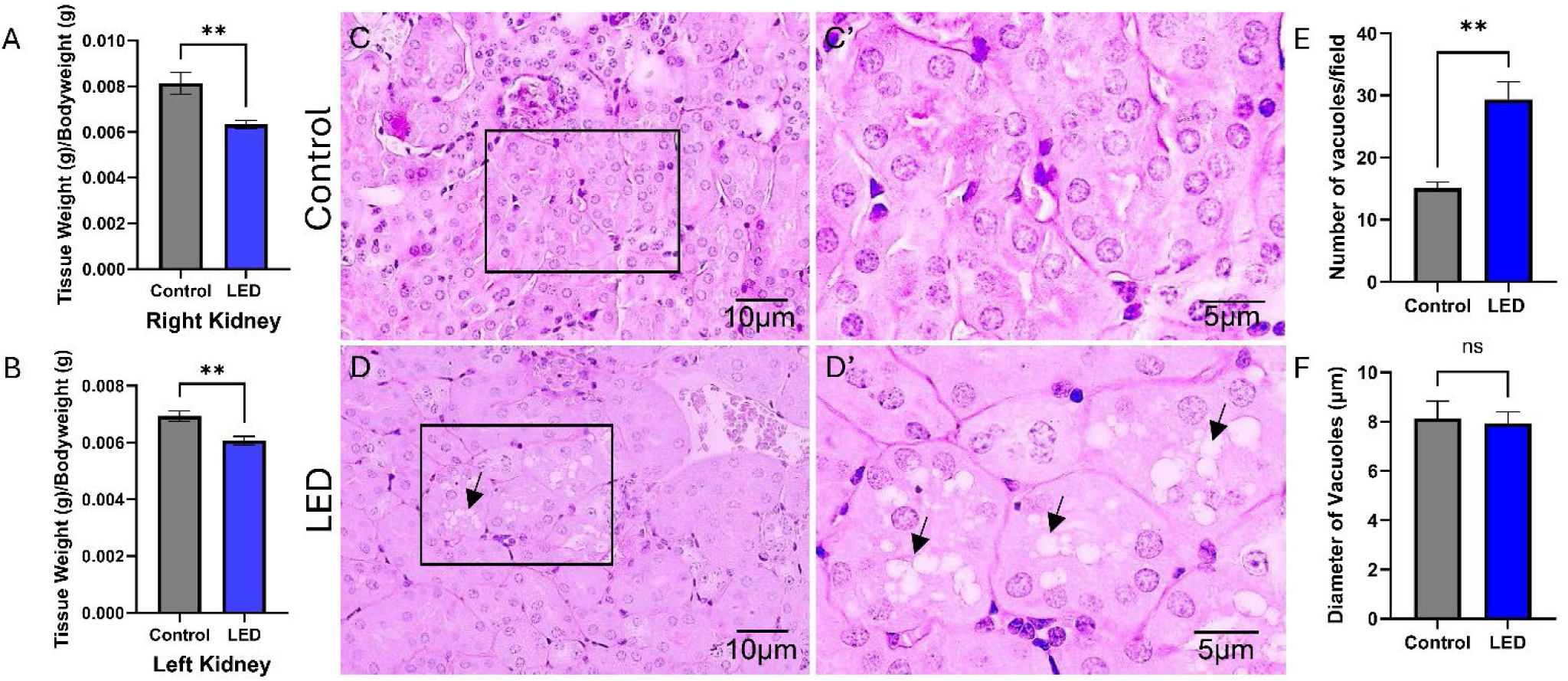
Renal tubular degeneration and vacuolation in mice exposed to 450 nm light. Histograms show normalized (A) right and (B) left kidney weight difference between control (grey) and 450 nm light-exposed animals (blue). Photomicrographs of kidney sections stained with Periodic acid–Schiff (PAS) stain from control (C) and 450 nm blue LED light-exposed mice (D). Regions indicated by rectangular boxes in C and D are shown at higher magnification on the right (C’ and D’). Note degenerating renal tubules with vacuolated cells (arrows) in blue LED light-exposed mice kidney. Bar graphs showing (E) average number of vacuoles per field and (F) average diameter of vacuoles in µm. Error bars indicate SEM. **p<0.01, n=10 per group for (A-B) and n=4 per group for (E-F). Data was analyzed by unpaired t-test.

Histological examination of kidney sections from control animals revealed normal tubular architecture with intact epithelial cells and minimal cytoplasmic vacuolation (Fig 4. C-C′). In contrast, kidneys from LED-exposed animals showed marked tubular epithelial degeneration characterized by extensive cytoplasmic vacuolation (arrows), cellular swelling, and disruption of normal tubular morphology (Fig 4. D-D′). Tubular epithelial vacuolation appeared predominantly focal, occurring in clusters of affected tubules rather than being uniformly distributed throughout the renal cortex and was absent or minimal in control animals. While no changes were observed in the diameter of vacuoles (p=0.8024; Fig 4. F), the total number of vacuoles per field significantly increased in kidneys (p<0.01; Fig 4. E) of LED-exposed animals.

Histological examination of liver sections from control animals revealed preserved hepatic architecture with normal hepatocyte morphology and limited cytoplasmic vacuolation (Fig 5. B–B′). In contrast, livers from LED-exposed animals showed marked hepatocellular vacuolation (arrows), cellular swelling, and disruption of normal hepatic architecture (Fig 5. C–C′). Oil Red O staining revealed minimal lipid accumulation in the livers of control animals (Fig 6. A–C). In contrast, livers from LED-exposed animals exhibited a marked increase in Oil Red O-positive lipid droplets throughout the hepatic parenchyma (Fig 7. D–F) indicating significant hepatic lipid accumulation and steatosis following chronic LED exposure. Morphometric analysis further supported this observation. Although the number of hepatic vacuoles was reduced (p<0.05; Fig 5. E) in the LED-exposed group compared with control animals, the mean vacuole diameter was significantly increased (p<0.0001; Fig 5. F), indicating the presence of fewer but substantially larger lipid vacuoles. This pattern is consistent with the progression of hepatic steatosis, in which enlargement and coalescence of lipid droplets result in the formation of larger intracellular vacuoles despite a reduction in their overall number.

**Figure 5.**
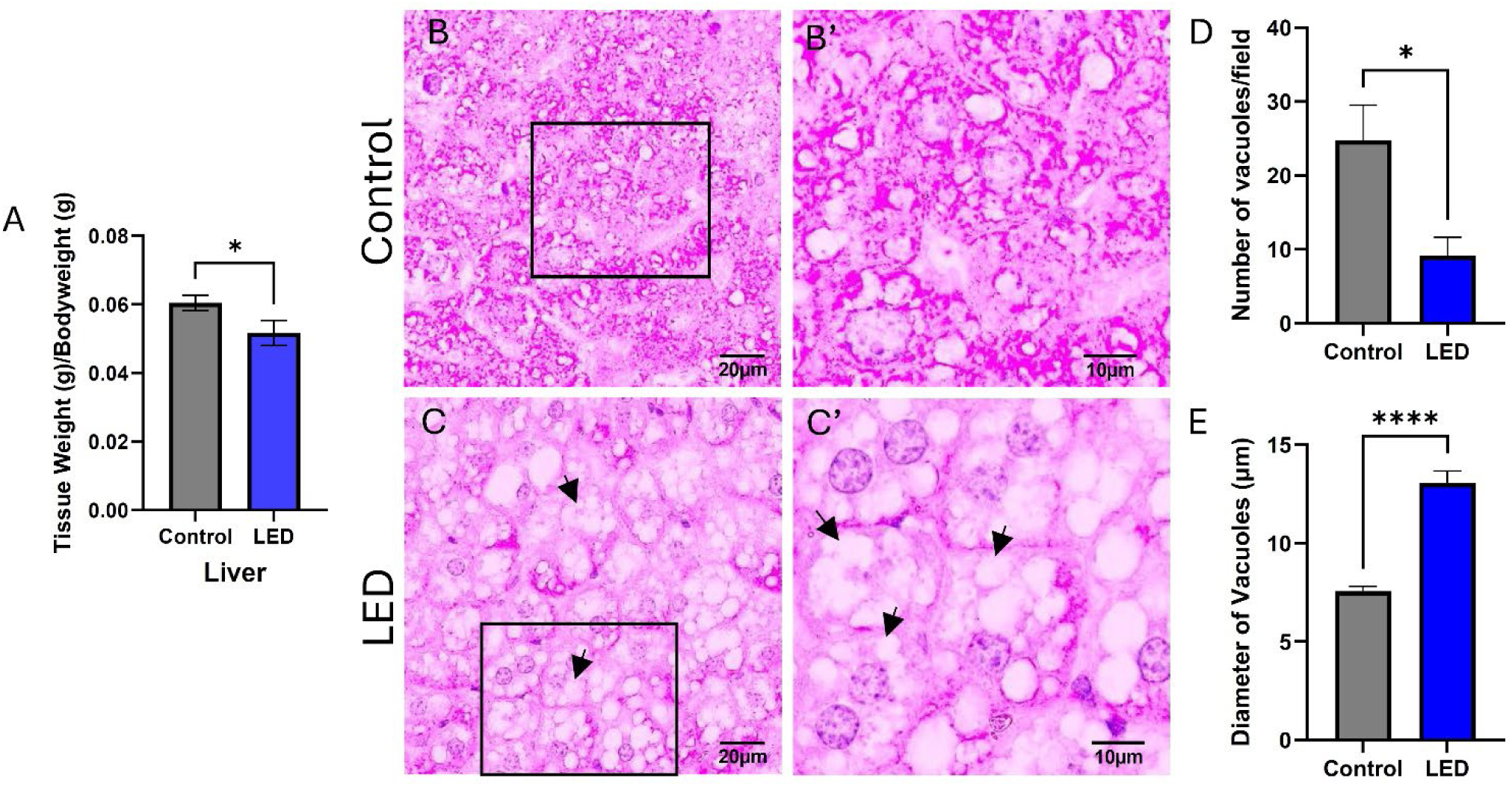
Hepatic vacuolation and tissue degeneration in mice exposed to 450 nm blue LED light. (A) Histogram shows a significant decrease in normalized liver weight in mice exposed to 450 nm LED light compared with controls. Photomicrographs of liver sections stained with Periodic acid–Schiff (PAS) stain from (B) control and (C) 450 nm blue LED light-exposed mice. Higher magnifications of regions indicated by rectangular boxes are shown on the right panel (B’ and C’). In the livers of blue LED light-exposed mice, hepatocytes increased in size, vacuolation (arrows) and widespread lipid accumulation, with peripherally displaced nuclei and decreased cytoplasmic glycogen when compared to control livers. Bar graphs showing (D) average number of vacuoles per field and (E) average diameter of vacuoles in µm. Error bars indicate SEM. *p<0.05, ****p<0.0001 n=10 per group for (A); n=7 per group for (B) and n=5 per group for (E-F). Data analysed by Student’s unpaired t-test.

**Figure 6.**
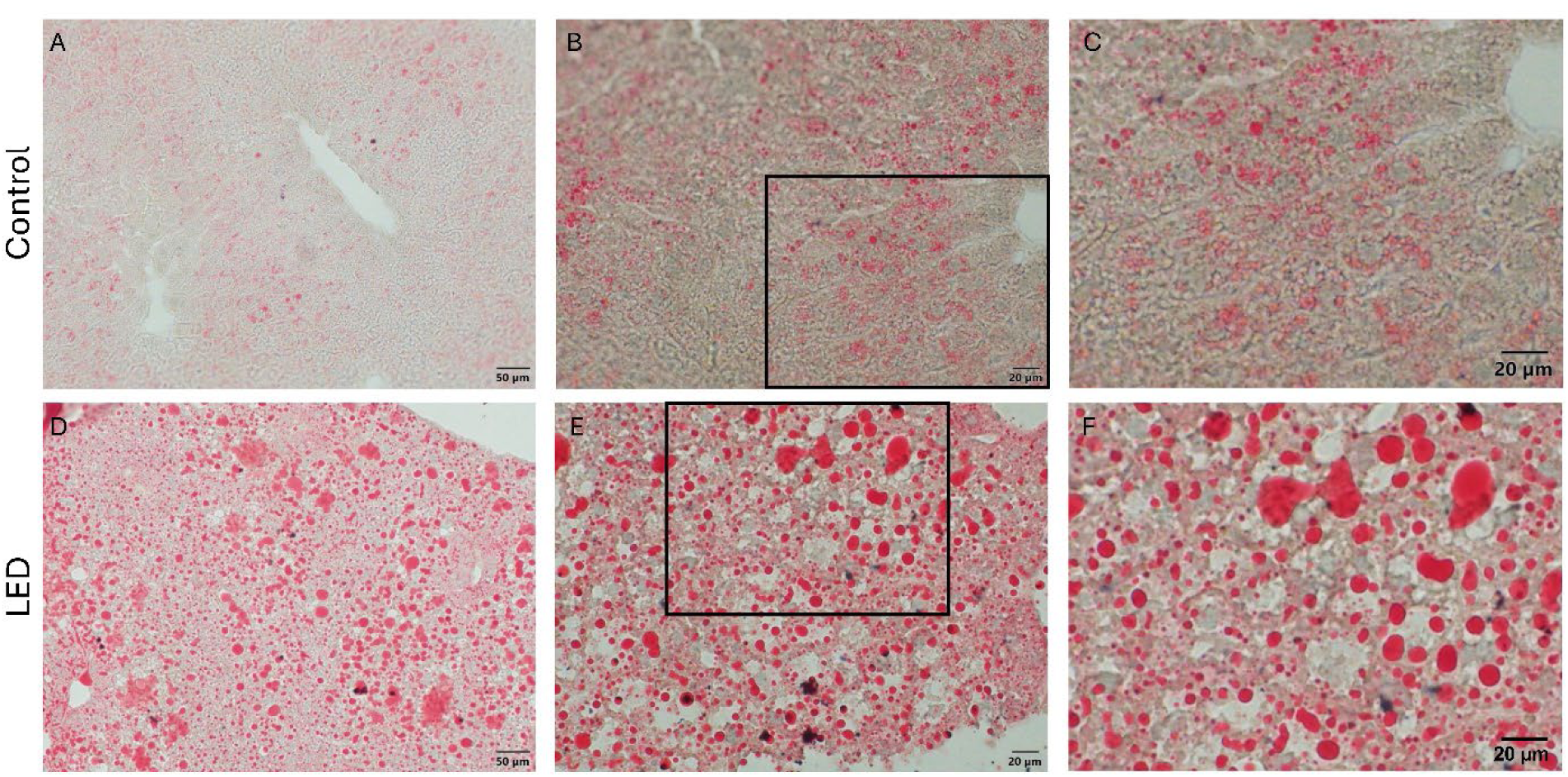
Liver. Accumulation of lipids in the liver of mice exposed to 450 nm. Mice exposed to 450 nm light accumulated excessive fat/lipid in their liver as revealed with oil red O staining. (A) Photomicrograph of liver sections from control (A, B, C) and 450 nm LED-exposed mice (D, E, F). Higher magnifications of the regions indicated by rectangular boxes are shown in C and F. Fat/lipid distribution in the liver of 450 nm exposed mice did not show any obvious distribution. Larger and smaller droplets were intermixed. While there were greater levels of staining in the 450 nm exposed animals, this was within the framework of reduced liver weight. Hence, organ reductions must have arisen from loss of normal liver tissue.

**Figure 7.**
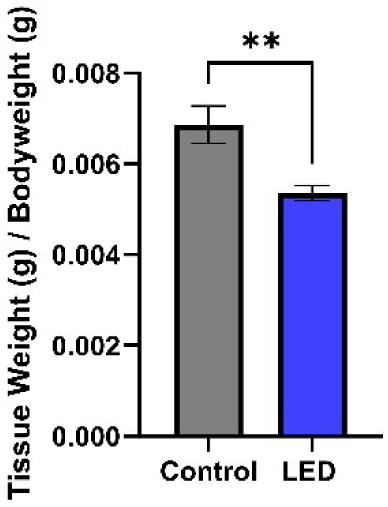
Heart. Mice exposed to 450 nm light show a reduction in total heart weight. Heart weight was normalized to body weight for analysis. The tissue appeared grossly normal. Tissues were not sectioned for further analysis because the appropriate plane of section was not obvious. Grey bars in the histograms are data from controls and blue from 450nm exposure Error bars indicate SEM. **p<0.01 n=10 per group.

Histological examination of testicular sections from control animals revealed normal seminiferous tubule architecture with a well-organized germinal epithelium containing abundant spermatogenic cells at different stages of maturation (Fig 8. A–B). Numerous mature spermatozoa were present within the lumen. In contrast, testes from LED-exposed animals exhibited disrupted seminiferous tubule organization, reduced numbers of mature spermatozoa within the lumen (Fig 8. C–D). Some spermatogenic cells appeared smaller with condensed nuclei, and increased intercellular spaces were observed within the germinal epithelium, indicating degeneration of spermatogenic cells and impaired spermatogenesis. Examination of sperm morphology demonstrated an increased incidence of abnormal sperm heads in LED-exposed animals compared with controls (Fig 9. B–D). The abnormalities included enlarged and distorted head shapes (arrows, Fig. D). Quantitative analysis showed a significant increase in the percentage of sperm exhibiting abnormal head morphology in the LED group (p < 0.05; Fig. B).

**Figure 8.**
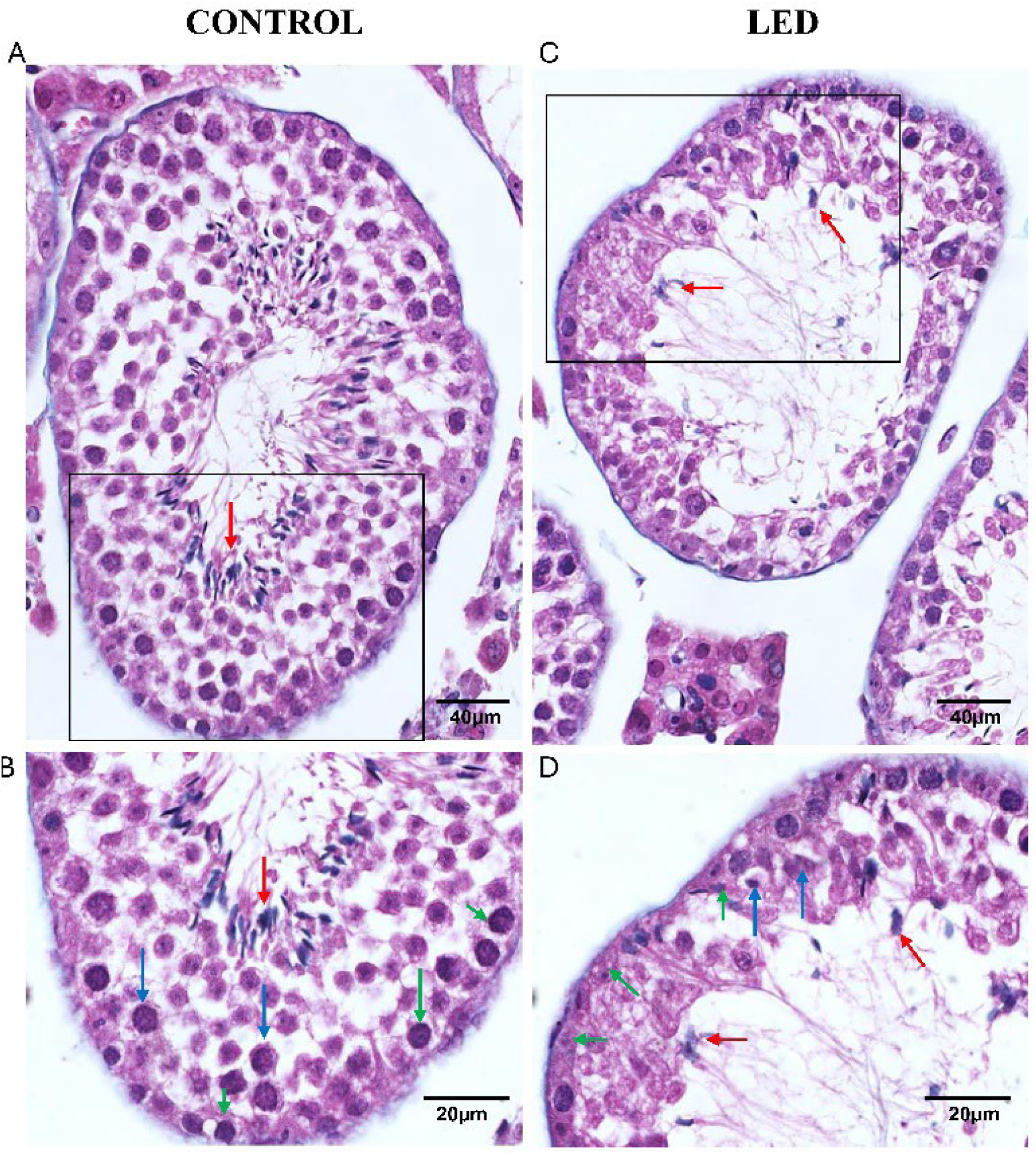
Testis. Abnormal morphology of testis in LED exposed mice. (A) Photomicrograph of testis sections showing seminiferous tubules from control (A and B) and 450nm LED exposed mice (C and D) stained with Masson’s trichrome stain. Regions indicated by rectangular boxes in the upper panel are shown at a higher magnification in the lower panel. Note less sperm in the lumen of the seminiferous tubule or attached to the Sertoli cells (red arrows) in LED-exposed group. In the control group spermatogonia are large, with prominent nuclei (green arrows) and primary spermatocytes and other cells are large with prominent nuclei (blue arrows). However, in LED group, spermatogonia are small with shrunken nuclei (green arrows) and primary spermatocytes and other cells are small with shrunken nuclei (blues arrows). Scale bar=40µm in upper panel and 20µm in lower panel.

**Figure 9.**
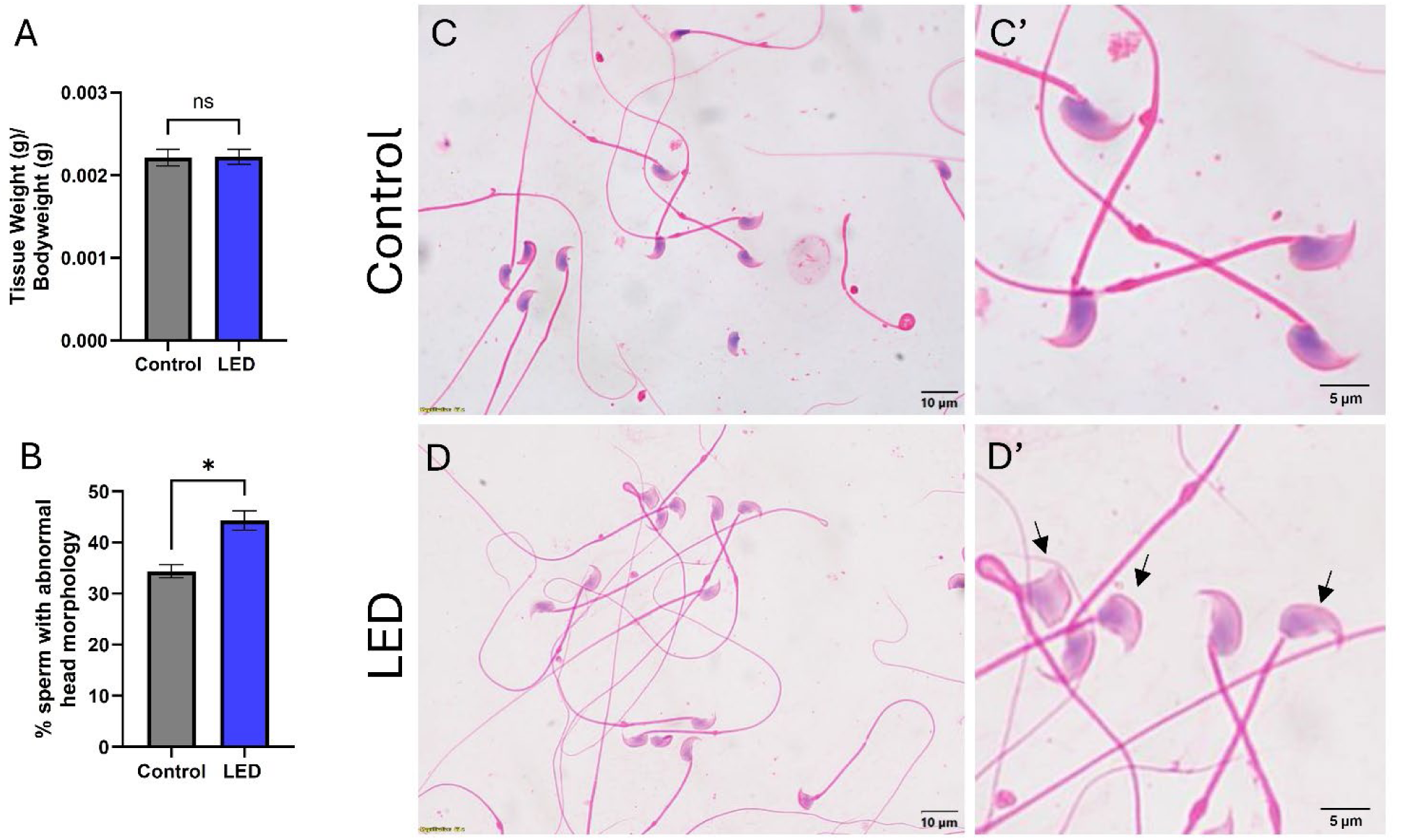
Sperm. Exposure to 450 nm blue light changes sperm head morphology. (A) Normalized testis tissue weights were the same for control and LED-exposed groups. (B) Percent sperm with abnormal head morphology where greater in LED-exposed mice. Photomicrographs of sperm smears stained with hematoxylin and eosin from control (C and C’) and 450 nm LED-exposed (D and D’) mice. Arrows indicate abnormal sperm heads in LED-exposed mice. Error bars indicate SEM. ns=not significant **p<0.01, n=3 per group; n≥162 sperms per animal. Data was analysed by unpaired t-test. Scale bar=10µm in C, D and 5µm in C’, D’

## Discussion

The present study demonstrates that long term exposure to narrow band 450 nm LED light was associated with increased body weight gain, greater white adipose tissue accumulation, altered glucose homeostasis, reduced relative organ weights, and histopathological changes in the liver, kidney, and testis. These changes occurred despite lower measured food intake in the monitored subset and in the absence of detectable changes in BAT weight or UCP1 protein expression. Together, these findings indicate that prolonged short wavelength LED exposure is associated with widespread metabolic and structural alterations in multiple organs.

Freshly excised mouse skin did not transmit detectable 450 nm light, consistent with its absorption by skin ^12, 13^. Hence, light did not impact directly on key organs, and their pathological changes are likely due to indirect mechanisms mediated, perhaps by the body surface and/or the eye. These findings suggest that direct irradiation of deep internal organs by 450 nm light was unlikely under the present experimental conditions. Consequently, the systemic metabolic alterations and histopathological changes observed in the exposed mice are more likely to involve indirect physiological pathways, although the underlying mechanisms were not investigated in the present study.

The 450 nm wavelength range is strongly absorbed by mitochondrial porphyrin ^14, 15^, raising the possibility that mitochondria within superficial tissues may participate in the initial biological response to short-wavelength light exposure. Previous studies have reported that 420 nm light exposure impairs mitochondrial function in Drosophila by reducing respiratory-chain activity and ATP production^16^. In mouse, retina real-time imaging reveals that 420 nm rapidly reduces mitochondrial respiration^15^. Hence, it is possible that there may be a mitochondrial involvement in a potential mechanism underlying the systemic effects observed in this study. We did not undertake any analysis of mitochondrial function in these mice; however other studies have clearly described a functional link.

Mice exposed to 450 nm LED showed a progressive increase in body weight despite consuming 10–15% less food than control animals, indicating a divergence between energy intake and weight gain. This finding suggests the possibility of altered metabolic regulation. However, additional studies incorporating direct measures of energy expenditure, metabolic rate and respiratory exchange ratio, will be required to determine the underlying mechanisms responsible for the increase in body weight. Post-mortem analysis of these animals revealed that weight gain was driven by selective expansion of white adipose tissue deposits with significant increases in both subcutaneous and epididymal fat depots. In contrast, neither brown adipose tissue weight nor UCP1 expression was altered by chronic 450 nm exposure. UCP1 is the principal mediator of brown adipose tissue thermogenesis and plays a critical role in energy dissipation through mitochondrial uncoupling ^17^. Therefore, the absence of changes in UCP1 expression demonstrates that the increased adiposity observed in LED exposed animals was not driven by a functional defect in classical brown fat thermogenesis ^17^. This suggests that the weight gain may be mediated by alternative pathways, such as altered peripheral insulin sensitivity ^18^.

Chronic 450 nm LED exposure resulted in a time dependent deterioration in systemic glucose homeostasis, with significant increases in OGTT and ITT AUCs from week 8 and peaking at week 20-32, indicative of progressive glucose intolerance and insulin resistance. This impairment likely correlates with the observed increase in total body mass, a physiological pattern highly consistent with compromised glucose metabolism observed in non-dietary models of metabolic dysfunction ^19^. Notably, these changes occurred despite largely normal fasting blood glucose levels. This pattern is consistent with an early metabolic disturbance, where alterations in glucose handling may be detected during physiological challenge before the development of sustained fasting hyperglycemia ^20^. Oral glucose tolerance testing is considered a more sensitive indicator of early metabolic dysfunction than fasting glucose measurements alone and may identify abnormalities in glucose regulation before overt diabetes develops ^21^.

By week 40, both OGTT and ITT AUCs declined relative to earlier time points, suggesting a partial metabolic adaptation despite continued exposure. This late-stage response may have resulted in a dissociation between basal and dynamic metabolic parameters, whereby fasting glucose levels normalized while glycaemic response during OGTT remained elevated. While the apparent normalization of ITT responses at week 40 suggests a reversal of systemic insulin resistance, the persistent impairment in OGTT indicates a divergence in glucose regulatory control. This pattern may reflect defects in pancreatic β-cell function, particularly impaired first phase insulin secretion, rather than ongoing peripheral insulin resistance^22, 23^, However, alternative mechanisms, including delayed insulin kinetics or tissue specific defects in glucose uptake, cannot be excluded. Notably, direct assessments of pancreatic function were not performed, limiting mechanistic resolution, but these findings provide a strong rationale for future investigation into light induced alterations in insulin secretory dynamics and β-cell workload ^24^. Mechanistically, the trajectory may reflect an initial disruption of circadian and metabolic regulation followed by compensatory adaptations, including peripheral clock recalibration, altered sympathetic signalling, and enhanced insulin secretion or sensitivity in the basal state. Light exposure is a key regulator of metabolic homeostasis through both circadian and non-circadian pathways, and chronic perturbation can initially impair insulin action while promoting longer-term adaptive remodelling of glucose regulatory systems^25^. Moreover, artificial light has been linked to circadian misalignment, insulin resistance, and increased diabetes risk, yet emerging evidence indicates that metabolic plasticity may partially mitigate these effects over prolonged exposure^26, 27^. Collectively, these findings suggest that chronic LED exposure drives early metabolic dysfunction followed by a phase of partial physiological adaptation, resulting in an IGT-like phenotype at later time points.

The reduction in the relative organ weight following chronic 450 nm exposure highlights the potentially systemic effect of this light exposure. The ∼10% organ shrinkage suggests impaired tissue maintenance or increased catabolic stress, a pattern compatible with the elevated cytokine shifts previously reported under 420-450 nm illumination^11^. Hepatic changes, including lipid accumulation and hepatocyte degeneration, are in line with previous reports linking short wavelength light exposure to altered liver metabolism^28, 29^. Similarly, renal tubular degeneration and testicular abnormalities observed in this study are consistent with patterns reported in experimental models of stress and metabolic disruption ^30, 31^. Importantly, the qualitative histopathological observations were supported by quantitative morphometric analyses. In the liver, chronic LED exposure produced marked neutral lipid accumulation demonstrated by increased Oil Red O staining together with significantly larger intracellular lipid vacuoles despite a reduction in vacuole number. This pattern is consistent with enlargement and coalescence of lipid droplets during the development of hepatic steatosis, a well-recognized morphological feature of fatty liver disease ^32, 33^. These observations are also consistent with recent reports demonstrating hepatic steatosis, inflammation, hepatocellular ballooning, and oxidative stress following chronic blue or LED light exposure^28, 29^. In the kidney, quantitative morphometry demonstrated a significant increase in tubular epithelial vacuole number without a corresponding increase in vacuole diameter, supporting the histological evidence of tubular epithelial injury. Similar renal tubular alterations have been described in experimental models of metabolic and oxidative stress ^30, 31^. Testicular vulnerability in the 450 nm LED cohort was equally marked, as evidenced by diminished sperm density, shrunken spermatogonia, and altered sperm head morphology. These distinct features are highly consistent with short wavelength LED induced reproductive injury. Because prior studies in male rats similarly note the degeneration of seminiferous tubules alongside impaired spermatogenesis^34^, our results reinforce the consensus that narrow band blue light acts as a potent driver of male gonad dysfunction. Collectively, these findings are consistent with the observation that chronic exposure to 450 nm LEDs produces a coordinated pattern of metabolic, hepatic, renal, and reproductive pathology.

Although this study was conducted in mice, the findings may provide insight into how altered light environments influence physiological processes. Modern LED lighting differs from natural sunlight in its narrower spectral range, particularly with increased relative short-wavelength content. In support of this, there are varied epidemiological studies revealing the importance of broad-spectrum sunlight on metabolism. Key among these is a comparison of 13,000 subjects that revealed improved insulin sensitivity in those exposed to sunlight^35^. The present results, together with existing evidence, suggest that such changes in light exposure may possibly be associated with alterations in human metabolism and tissue function. However, differences in species biology, exposure conditions, and environmental context limit direct translation to humans. Further studies are required to clarify the relevance of these findings to human health.

## Methods

### Animals and Housing

Male C57BL/6J mice (stock no. 000664, Jackson Laboratory, Bar Harbor, ME, USA) were maintained under a 12:12 h light–dark cycle with controlled temperature and humidity. At three months of age, mice were allocated to control or blue-light–exposed groups (n = 14 per group; total n = 28). Food and water were provided *ad libitum* throughout the study. For a randomly selected subset of animals (n = 5 per group), food intake was assessed weekly. Briefly, 100 g of standard chow was provided to each singly housed animal at the beginning of each week, and the remaining food was weighed after 7 days. Weekly food consumption per animal was calculated as the difference between the initial and remaining amounts and was normalized to body weight. Cumulative food intake per animal over the 40-week study period, and average weekly food intake normalized to body weight, were plotted. Body weight and fasting blood glucose levels were measured bi-weekly in all animals. Animal rooms were illuminated with three identical Osram L36W/76 daylight fluorescent strip lighting in each room to ensure consistent lighting. Control and experimental animals were housed in separate rooms under identical background lighting, temperature, and diet. Animal care and treatment protocols adhered to the guidelines approved by the Institutional Animal Care and Use Committee of Kuwait University, and in compliance with the NIH Guidelines and the Guide for the Care and Use of Laboratory Animals (Approval No. 23/VDR/EC/3560).

### Short-Wavelength Light Exposure

Our previous work^11^ showed that the physiological impact of short wavelength LED light exposure is largely similar at 420 and 450 nm. Hence, here we focused on 450 nm LEDs. Control animals were housed under standard fluorescent lighting (Osram L36W/76), which emits a broad-spectrum white light distinct from the narrow-band 450 nm LED used in experimental conditions. Experimental animals were exposed to this LED light generated by 24 regularly spaced diodes mounted on an aluminium PCB (10.7 × 8.2 cm) positioned above the cage lids. Each LED was equipped with a 90° lens, and the emission had a half-power bandwidth of ∼14 nm. Irradiance at cage floor level was 13 mW/cm², as measured with a calibrated radiometer. This is comparable to human exposures in the built environment. Mice were exposed to the LEDs for 5 h per day, from 11:00 to 16:00, within the light period of a 12:12 h light–dark cycle, for the 40 weeks duration of the study, while controls were not exposed to LED lighting.

### Light penetration

To determine whether 450 nm light penetrated beyond the skin, a single mouse was euthanized by cervical dislocation and skinned. The skin was placed with its outer (hair) side directly against the 450 nm LED array in a dark room. An ILT5000 radiometer (International Light Technologies) and an Ocean Optics QE65 Pro spectrometer were positioned on the inner (dermal) side to detect transmitted light.

### Metabolic Tests

Oral glucose tolerance test (OGTT) and insulin tolerance test (ITT) were performed as previously described^36^. Briefly, mice were fasted for 6 hours, and body weights were recorded. Blood glucose levels were measured from the tail vein blood using a glucometer (One Touch Select, China). Each mouse was orally administered 2g/kg glucose for OGTT using a gavage needle. For ITT, each mouse was injected intraperitoneally with 0.5IU/kg of insulin. Blood glucose levels were recorded at 0 min (before glucose/insulin administration) and at 15, 30, 60, 90 and 120 min post-administration. Metabolic tests were performed before LED exposure (week 0) and at weeks 4, 8, 20, 32 and 40 post-exposures.

### Tissue Collection

Mice were sacrificed by cervical dislocation, and the following tissues were harvested: subcutaneous white adipose tissue (SWAT), epididymal white adipose tissue (EWAT), brown adipose tissue (BAT), liver, heart and kidneys. An incision was made along the abdominal skin to collect SWAT (Inguinal) from the posterior end of the body. EWAT was collected by opening the peritoneal cavity, and BAT was collected by exposing the fat deposits below the nape. The heart, liver, both kidneys, and the testes were collected in ice cold PBS, cleaned to remove any hair or other contaminants, gently rinsed, and weighed. Tissues were fixed in 4% paraformaldehyde for histopathology or snap-frozen for western blotting. The cauda epididymis was collected in PBS (pre-warmed to 35°C) and gently minced to release the spermatozoa.

### Western Blot Analysis

Frozen BAT was lysed in RIPA buffer with protease and phosphatase inhibitors (Chemcruz, sc-24948), using a Bullet Blender tissue homogenizer (Next Advance). Samples were centrifuged to separate the fat layer, and the lysate was used for further analysis. Protein samples were resolved on 4–20% Mini-PROTEAN® TGX Stain-Free™ protein gels (Bio-Rad, 4568096) and transferred to PVDF membranes (Bio-Rad, 1704156) using the Bio-Rad Trans-Blot Turbo transfer system. Membranes were blocked in 1% casein (Bio-Rad, 1610782) and incubated with the following primary antibodies: anti-UCP1 (Abcam, ab10983), and anti-GAPDH (Cell Signalling Technology, #2118), followed by a goat anti-rabbit HRP secondary antibody (Vector Laboratories, PI-1000-1). Anti-UCP1 (ab10983) has been demonstrated to work efficiently for western blots on proteins from brown adipose tissue^37^. Blots were developed using Clarity western blot substrate (Bio-Rad, 170-5061) on ChemiDoc System (Bio-Rad). Image Lab software was used to quantify band intensities.

### Histopathology Staining

Fixed liver, kidney and testes were dehydrated and cleared using a Leica automated tissue processor. Samples were embedded in paraffin, and 5 μm thick sections were cut on a microtome (Leica). Periodic Acid-Schiff (PAS) staining was performed using Schiff’s reagent and hematoxylin. Hematoxylin and Eosin (H&E) staining was performed using standard histological procedures. To quantify vacuole number and diameter, 5-6 images were obtained from each animal and ImageJ was used for quantification. For Oil Red O staining, fixed frozen livers were sectioned at 10 μm on a cryostat. These were stained with Oil Red O to visualize lipid, and counterstained with hematoxylin.

A 10μl suspension containing spermatozoa was spread on a slide’s surface and air dried. Smears were fixed in 95% ethanol for 15 minutes, followed by staining with hematoxylin and eosin. Slides were air dried and mounted using DPX. Sperm heads were characterized as abnormal if the head hook was absent, the head was detached from the tail (tailless), or the head was shaped banana-like or amorphous^38^.

### Quantification and Statistical analysis

Data were analysed using GraphPad Prism (GraphPad Software, San Diego, CA, USA) and are presented as mean ± SEM. Student’s *t*-test was used to compare two groups when data were normally distributed. For OGTT and ITT, the area under the curve (AUC) for blood glucose over time was calculated and compared between groups by Student’s t-test. A p-value < 0.05 was considered statistically significant. Statistical details, including the number of animals and samples, statistical tests used, and levels of significance are indicated in the figure legends.

## Data Availability

All relevant data are reported in this manuscript. This paper does not report original code. Any additional information required to reanalyze the data reported in this paper is available upon request.

## Acknowledgements

We thank the Animal Resources Center (Kuwait University) staff for technical support. The authors would like to thank Dr. Josely George, Maria Joji and Ashley Josely.

## Author contributions statement

Conceptualization, H.A-H.; methodology, H.A-H., L.D., S.H., M.A-O., and S.M; Investigation, H.A-H., L.D, S.H., M.A-O., R.K, B.S., S.R., B.A.; writing—original draft, H.A-H., L.D., and G.J.; writing—review & editing, S.H., M.A-O., and S.M.; funding acquisition, H.A-H.; resources, H.A-H.;supervision, H.A-H., S.H., G.J. All authors reviewed manuscript.

---

**Supplementary Figure 1.**
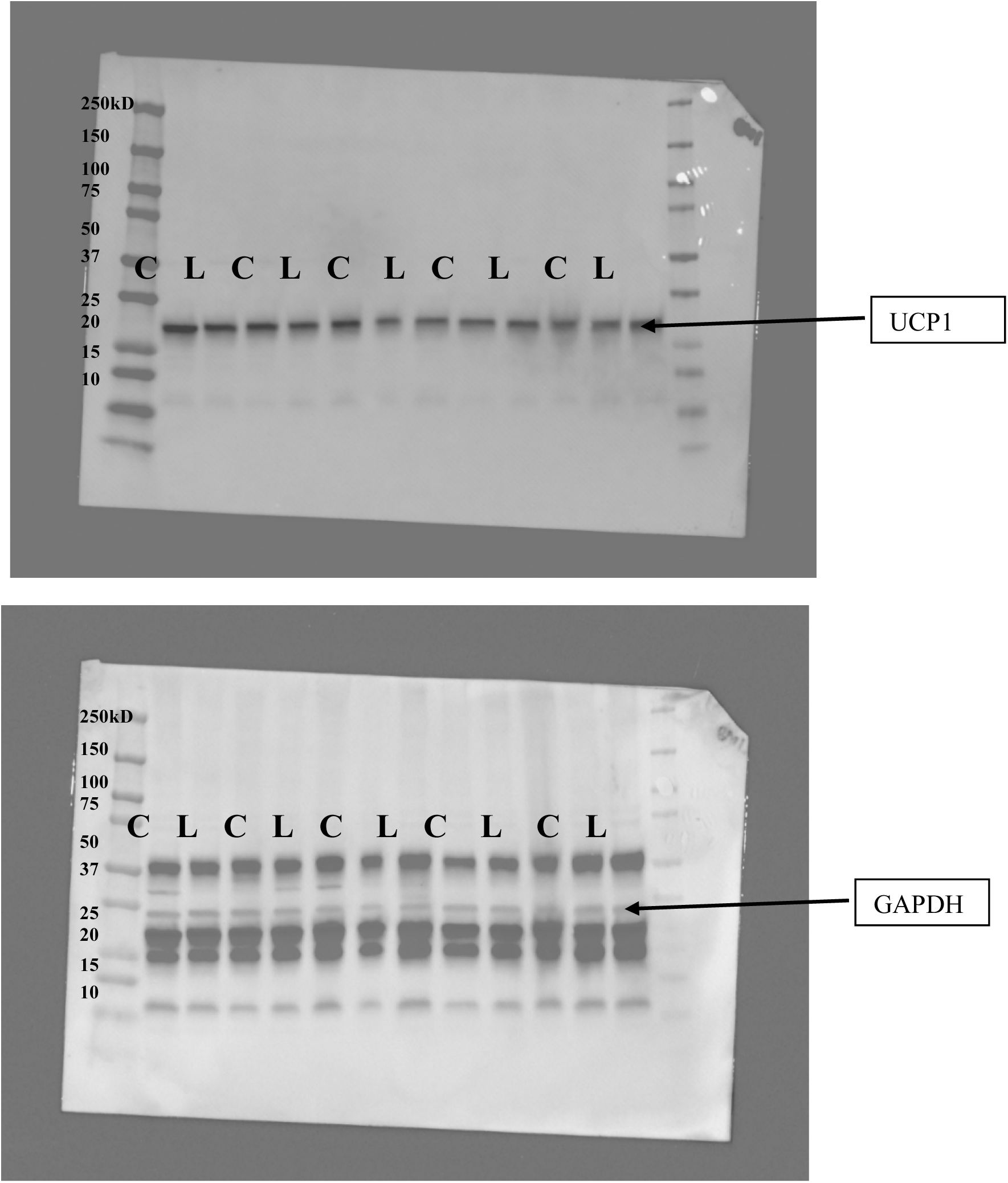
Original Western Blots for UCP1 and GAPDH. Arrow indicates band of interest.

